# Rapid *Agrobacterium*-mediated transformation of tobacco cotyledons using toothpicks

**DOI:** 10.1101/204891

**Authors:** Yuan-Yeu Yau, Mona Easterling, Lindsey Brennan

## Abstract

Tobacco is a model plant for genetic transformation, with leaf-disk transformation being the most commonly used method for its transformation. One disadvantage of leaf-disk transformation is obtaining an adequately sized leaf. Cotyledons from young seedlings are considered too small and fragile to use. In an attempt to overcome this drawback, a protocol was developed using toothpicks as a tool to inoculate cotyledons ~2mm in diameter. *Agrobacterium tumefaciens* LBA4404 hosting two different plasmids (pC35.BNK.2 or pRB140-Bxb1-op) was used for transformation. Fifty-six putative transgenic shoots (T_0_) were obtained from pC35.BNK.2 transformation. Among them, 38 (68%) grew roots in kanamycin-containing medium. Approximately, 35% of transgenic lines contained a single-copy transgenic locus based on Mendelian inheritance analysis and chi-square (χ^2^) test of T_1_ seedlings from 17 lines. To simplify the protocol, water-prepared *Agrobacterium* inoculum was used in pRB140-Bxb1-op (containing *gus* gene) transformation. This resulted in ~35% putative T_0_ transgenic lines stained strong blue with GUS histochemical staining assay. Both sets of results demonstrate toothpick inoculation to be an effective approach for *Agrobacterium*-mediated tobacco cotyledon transformation. This reduces wait time required in existing leaf-disk transformation method using mature leaves. Removal of step requiring submersion of explants in *Agrobacterium* liquid culture, the protocol also has advantages by minimizing *Agrobacterium* overgrowth and maintaining explant fitness for later tissue-culturing.

## Introduction

Tobacco is a model plant for *Agrobacterium*-mediated genetic transformation due to the simplicity of its transformation procedures. The traditional technique does not require expensive machinery or complicated procedures. Some significant advantages of using tobacco for genetic transformation are: (1) Tobacco plants can be easily regenerated from tobacco leaf pieces through organogenesis (Constantin et al. 1977), (2) Short acclimatization time and a high transpotting survival rate: up to 100% of the *in vitro*-raised plants transferred from lab to greenhouse condition were successfully established *ex vitro*. The acclimatization is brief, taking only a matter of days (Chandra et al. 2010), (3) Ability to maintain a hemizygous state with T-DNA cassette transfer: the inserted T-DNA cassette can be maintained at hemizygous status in a sterile environment through simple vegetative propagation. This can be achieved through cutting of tips or stems and reproduction in solid MS medium (Murashige and Skoog 1962) without phytohormone. The simplicity of the hemizygous T-DNA cassette is necessitated in some research projects. For example, hemizygous T-DNA structure has been used for site-specific deletion or integration experiments using site-specific recombinases (Hou et al. 2014), (4) Easy crossing: due to large flower size, hand-pollination is easily accomplished, (5) Longevity: by removing the flowering buds or tips, plants continue growing in greenhouse conditions for extended periods of time, which provides supplemental experimental material, particularly for the W38 tobacco species, (6) Prolific seed production for sustaining lines and testing results, (7) Increased biomass with the potential for molecular farming to produce recombinant proteins, due to tobacco’s high biomass yield (Twyman et al. 2003), (8) A model plant for agoinfiltration transient assays (Ma et al. 2012). Due to the variety of advantages mentioned above, most plant research scientists view tobacco as a prime choice for genetic transformation in proof-of-concept experiments with the added benefit of multiple, practical uses.

Currently, leaf-disk transformation is the most frequent used method for tobacco genetic transformation (Horsch et al. 1985). However, by using this method, an adequate size of *true leaf* (as oppose to cotyledons) tissue is required for cutting into leaf disks (usually ~ <u>1cm</u> in diameter). It is not practical to use very young tobacco cotyledons, for example 10-day-old cotyledons (~ <u>2 mm</u> in diameter), as an experimental material. The tiny size and fragility of cotyledons limits the use of forceps for manipulation, and it is difficult to handle such tiny tissue disks in transformation solution as well. Therefore, waiting for the true leaves to reach a bigger size before transformation use is necessary. This process can take a couple of months from seeds to use. In addition, in a re-transformation project (performing second transformation on a transgenic line), the seeds of the first-transformed lines needed to be selected on an antibiotic-containing medium beforehand, to ensure the presence of the first transgenic cassette. During the process of selection, the growth of surviving (antibiotic-resistant) seedlings is generally slow and stunted. This is especially true when the selection agent is hygromycin, as hygromycin is generally more toxic than kanamycin. To prevent the loss of transgenic plants, seedlings are usually transferred to fresh medium which does not contain antibiotics to stabilize their growth following selection. Using true leaves, the wait can usually be measured in weeks for plant material to be sufficient in size for use in a second transformation. Using cotyledons of the surviving seedlings following selection for second-run transformation, would save weeks of time and funds. Therefore, it is desirable to develop a method for transforming cotyledons (instead of true leaves) from the earliest stage of tobacco seedlings.

In this study, we explore the possibility of using cotyledons from young tobacco seedlings for transformation by using sterile toothpicks to deliver the *Agrobacterium* for infection. Two experiments were conducted: (1) Re-transformed cotyledons of transgenic tobacco lines (previously transformed with binary vector, pN6.Bxb1, which contains a **hygromycin**-resistant gene) with another binary vector (pC35.BNK.2) containing **kanamycin**-resistant gene, and (2) repeat procedures mentioned in (1) again with another plasmid pRB140-Bxb1-op (with a *gus* gene), using *Agrobacterium* prepared in water (Fig. 1A). The final transgenic lines should contain both T-DNA cassettes and both hygromycin- and kanamycin-resistant genes.

**Fig. 1.**
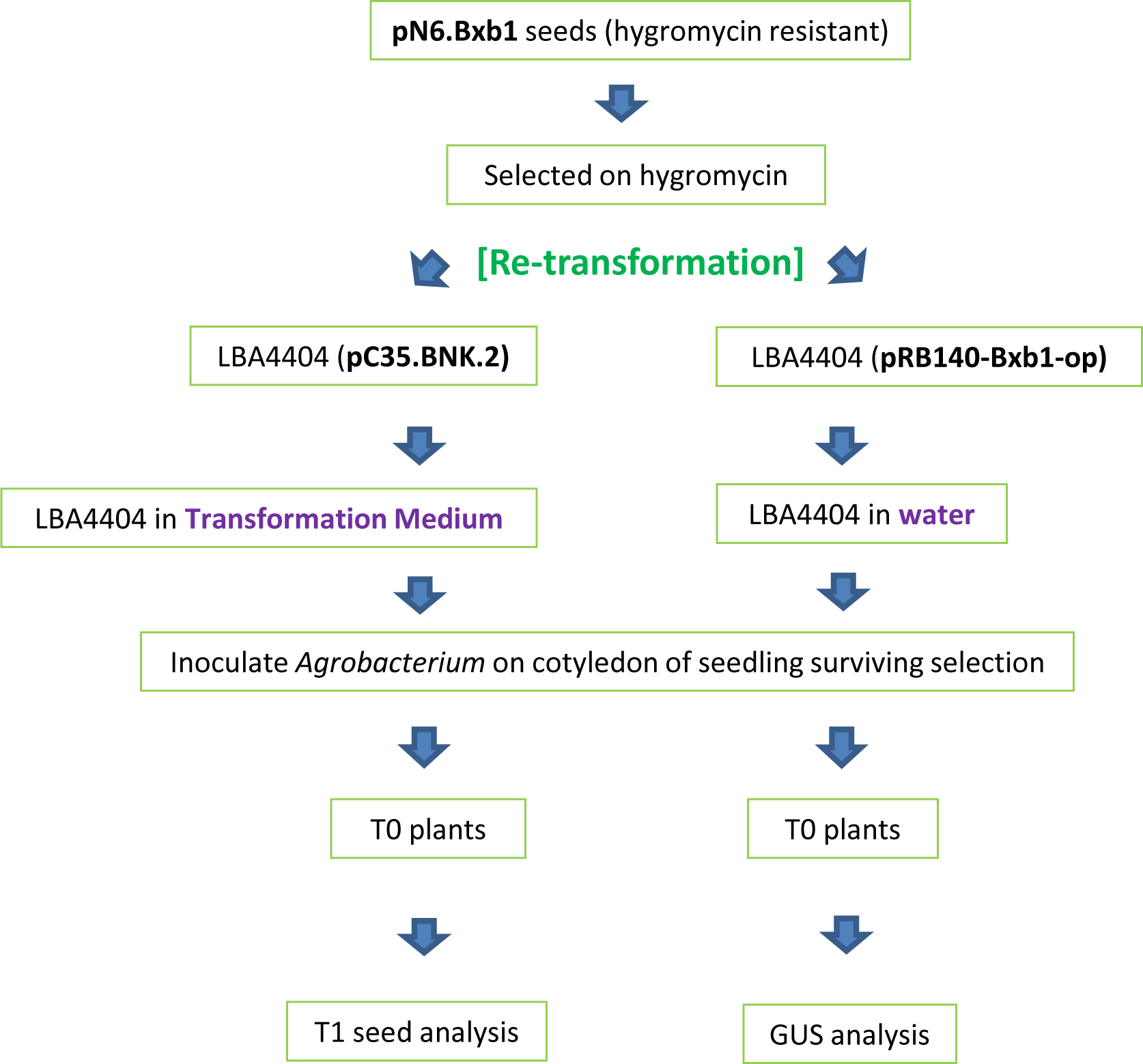
Outline of genetic transformation of this study.

## Materials and methods

### Plant materials and tissue-culture conditions

Seeds of transgenic tobacco (*Nicotiana tabacum* L. cultivar “Petit Havana” SR1) pN6.Bxb1 were germinated. These T_1_ seeds were from a previous project (Thomson et al. 2012). The project was functional study of site-specific recombination system Bxb1-*att* in plants. Seeds were sterilized with 70% ethanol for 2 min and bleach (sodium hypochloride) [30% (v/v), and drops of Triton-X 100] for 20 min, and washed thoroughly with autoclaved distilled water. Sterilized seeds were germinated on MS medium, which contains MS mineral salts (Murashige and Skoog 1962; Cat. No. M524, *Phyto*Technology Lab), 3% (w/v) sucrose, 1x Gamborg’s vitamin solution (Cat. No. G1019, Sigma-Aldrich), 0.8% agar and 45 µg/mL hygromycin (Cat. # H3274, Sigma, USA). Plates were sealed with a medical air-permeable tape (Micropore™ Surgical Tape; 3M Health Care, USA) (Clarke et al. 1992) and placed in a 25°C growth chamber with 16-h/8-h (light/dark) photoperiod. Seedlings that displayed stunted growth, a pale green to yellowish cotyledons, and inhibition of hypocotyl extension were considered susceptible to hygromycin. Seedlings with healthy green cotyledons and roots are considered hygromycin-resistance. Cotyledons of survived seedlings were used for genetic transformation.

### *Agrobacterium* strain and binary vectors

Detailed procedures for constructing pC35.BNK.2 were described earlier (Yau et al., 2011). Construct of binary vector pRB140-Bxb1-op (**Fig.1B**) was also described previously (Yau et al., 2012). pC35.BNK.2 uses pCambia2300 as a backbone, which carries the *nptII* gene (confers kanamycin resistance). *Agrobacterium tumefaciens* LBA4404 was used to host either pC35.BNK.2 or pRB140-Bxb1-op for genetic transformation. The vectors were electroporated into ElectroMax™ *Agrobacterium tumefaciens* LBA4404 competent cells (Cat. No. 18313-015, Invitrogen, USA) separately using an electroporator (Multiporator®, Eppendorf). Forty µL LBA4404 competent cells and 3 µL plasmid were mixed and then transferred into a 1 mm-gap electroporation cuvette (Cat. No. 94000100-5, Eppendorf, USA). The electroporation was carried out using a manufacturer pre-loaded program designated for bacterial electroporation (2000V, Time constant: 5.0 milliseconds). One ml LB liquid medium was then added into the electroporation cuvette and mixed with the electroporated competent cells. The mixture was then transferred to a Falcon^®^ 14-ml polypropylene round-bottom tube (Becton Dickinson Labware, USA) and incubated at 30°C for 3 hours with a 225-rpm shaking.

After 3 hours, the bacterial culture of 20 µl, 50 µl or 100 µl was spread onto LB + streptomycin (100 µg/ml) + kanamycin (50 µg/ml) plates. The streptomycin was used to select *A. tumefaciens* LBA4404 cells’ disarmed Ti pAL4404, and kanamycin was used to select transformed bacteria. Plates were placed in a 30°C incubator for 2-3 days to produce colonies.

### Prepare *Agrobacterium* for transformation

For first experiment, single colonies (derived from pC35.BNK.2 electroporation) from plates were picked with a 15-cm sterile Cotton Tipped Applicator (Puritan Medical Products Company, Guilford, Maine, USA) and streaked on LB plates containing antibiotics streptomycin and kanamycin, and allowed to grow at 30°C for 1 day. For tobacco genetic transformation, *Agrobacterium* grown overnight was scraped from the plates with a sterile inoculation loop and suspended in 100 µL Transformation Medium [MS mineral salts, 3% (w/v) sucrose, 1x Gamborg’s vitamin solution, 3 µg/mL 6-Benzylaminopurine hydrochloride (Cat. No. B5920, Sigma-Aldrich) and 100 µM Acetosyringone (AS) (Cat. No. D134406, Sigma-Aldrich)]. Transformation Medium was adjusted to pH5.8 with 0.1N KOH or HCl and autoclaved at 121°C and 120 Kpa (1 PSI = 6.89 Kpa) for 20 minutes. AS was dissolved in 70% ethanol and added to the cooled autoclaved medium (Jones et al 2005). AS is a phenolic compound that stimulates the induction of *Agrobacterium* virulence genes and improve the transformation efficiency (Nadolska-Orczyk and Orczyk 2000). *Agrobacterium* colonies were suspended in Transformation Medium, and were then diluted 10 times with the same medium for genetic transformation. For the second experiment, colonies (derived from pRB140-Bxb1-op electroporation) was directly picked and dissolved in autoclaved water (not Transformation Medium) for toothpick inoculation.

### *Agrobacterium-*mediated transformation

The process of using sterile toothpicks to inoculate *Agrobacterium* to tobacco cotyledons was summarized in diagram Fig. 2. Two-week-old seedlings surviving hygromycin-selection were pulled out from plates and placed in another sterile plate to cut off the roots in an ESCO Horizontal Airstream^®^ Laminar Flow hood (ESCO, USA) (Fig. 2A and 2B). Cotyledons, ~ 2.5mm in diameter, were gently bruised near the center with a sterilized (autoclaved) point-ended toothpick. The toothpick had dipped into the *Agrobacterium* (containing pC35.BNK.2) suspension described above (Fig. 2C). After inoculation, the cotyledons (on the leftover stem) were placed abaxial on co-cultivation medium [Transformation Medium solidified with 0.8% agar (Cat. No. A7921, Sigma)] for 3 days in dark, and then transferred to selection medium [Transformation Medium + cefotaxime/carbenicillin (500 µg/mL) + kanamycin (100 µg/mL), and solidified with 0.8% agar], with the leftover stem sticking into the medium (Fig. 2D). Mixture of 50% (w/w) cefotaxime (Cat. No. C380, *Phyto*technology Lab., USA) and 50% (w/w) carbenicillin (Cat. No. C346, *Phyto*technology Lab., USA) were used together to remove *Agrobacterium*. Plates were sealed with an air-exchangeable 3M Micropore™ tape and placed in a growth chamber with 16-hr light/8-hr dark photoperiod. Sub-culturing was carried out every two weeks. For pRB140-Bxb1-op transformation, the procedures were similar to that of pC35.BNK.2 transformation, but *Agrobacterium* solution was prepared by simply suspending bacterial colonies in autoclaved water (not Transformation Medium). pRB140-Bxb1-op contains a GUS gene. GUS expression in putative transgenic plants was evaluated and documented.

**Fig. 2.**
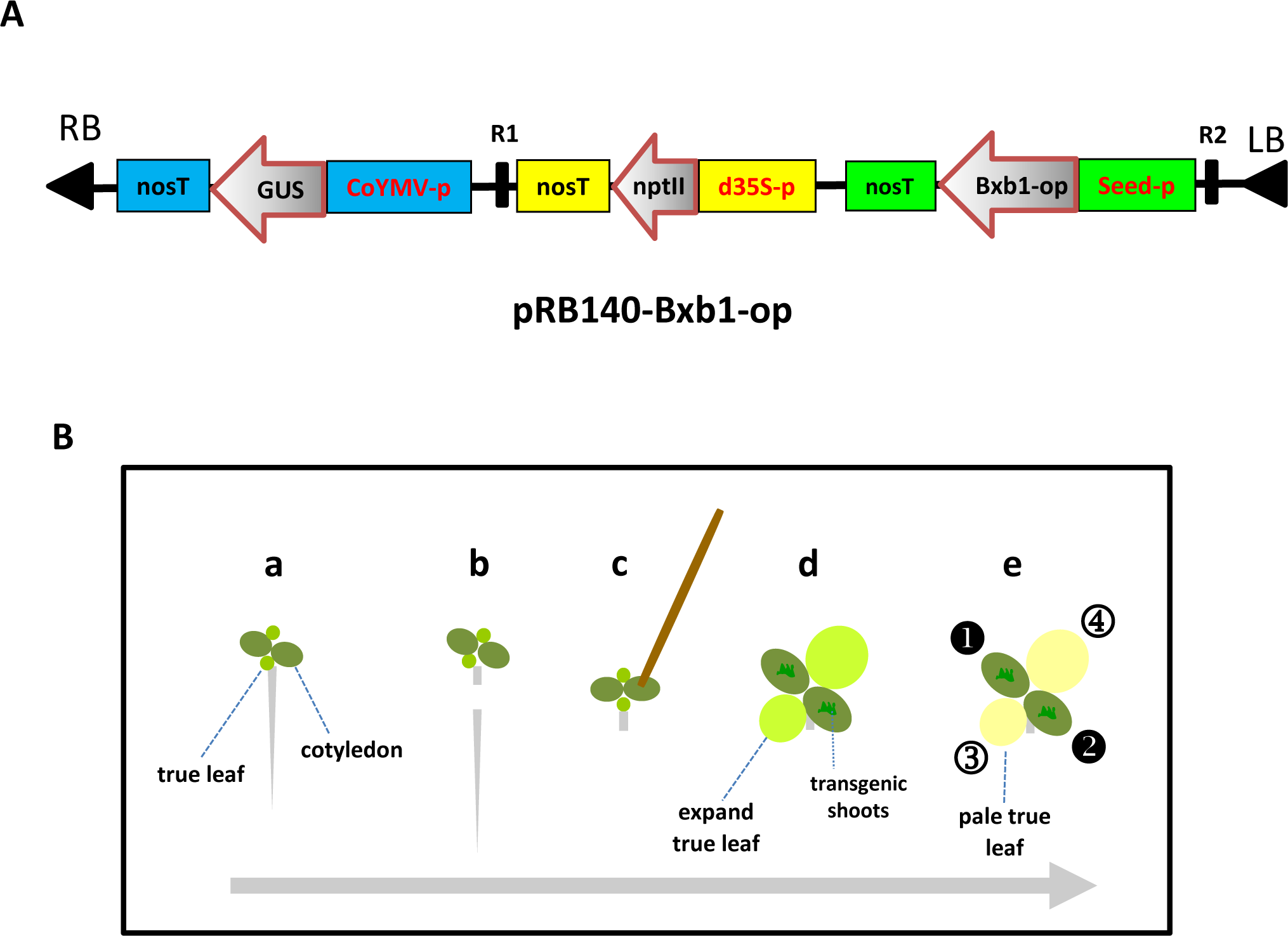
A Linear (partial) schematic cassette of binary vector pRB140-Bxb1-op T-DNA. LB/RB: left/right border of *Agrobacterium*, nosT: nopaline synthase (NOS) terminator, GUS: β-Glucuronidase gene, CoYMV-p: Commelina yellow mottle virus promoter, nptII: neomycin phosphotransferase II gene, d35S-p: double cauliflower mosaic virus 35S promoter, Bxb1-op: codon-optimized site-specific recombinase gene *bxb1*, seed-p: seed specific promoter. R1/R2: Bxb1-*att* site-specific recombination system recognition sites. **B** Schematic representation of *Agrobacterium* inoculation of tobacco cotyledons with sterile toothpicks for genetic transformation. **a** Two week-old seedlings with two large cotyledons (dark green oval) and two small true leaves (light green oval). **b** Cut root, a small portion of the root still remaining with the two cotyledons and two true leaves. **c** Inoculation of *Agrobacterium* with a sterile toothpick. **d** Size of both cotyledons and true leaves expand quickly. Putative transgenic shoots appear from the two *Agrobacterium*-infected cotyledons. **e** Putative transgenic shoots grow and true leaves (circles 3 and 4) become yellowish after antibiotic selection.

### Kanamycin selection of putative transformants from secondary transformation

Two weeks into kanamycin selection, the transformed cotyledons were separated from the stem with a sterilized scalpel and placed on freshly-prepared selection (kanamycin) medium. Putative transgenic shoots from the bruised region of the cotyledons were allowed grow further. Shoots 1-cm in length were cut and transferred into Rooting medium [MS mineral salts, 3% (w/v) sucrose, 1x Gamborg’s vitamin solution, 0.8% agar] supplemented with 100 µg/mL kanamycin and 400 µg/mL of mixture of cefotaxime and carbenicillin. One to two putative transgenic shoots were excised from every single cotyledon. Rooted plants were allowed to grow to 5-cm in a Magenta^®^ Plant Tissue boxes, and then transferred to soil.

### Genomic DNA extraction

A portion (a 1/4-size cap of a 1.5-ml microcentrifuge tube) of each leaf excised from putative transgenic plants or controls in the Magenta^®^ boxes was harvested into 1.5-ml microcentrifuge tubes. 400 µL grinding buffer (200 mM Tris-HCl, pH5.7, 250 mM NaCl, 25 mM EDTA and 0.5% SDS) was added to each tube and ground with a Kontes pellet pestle^®^ (VWR, Batavia, IL, USA) driven by an overhead stirrer (Cat. No. 2572101, IKA Works Inc., USA). The ground samples were centrifuged for 5 minutes at maximum speed (16,800 x *g*) with an Eppendorf benchtop centrifuge (Centrifuge model 5418). 300 µL of supernatant was transferred to a new microcentrifuge tube and 300 µg/mL of isopropanol was added to precipitate genomic DNA. After inverting several times, mixture was centrifuged for an additional 15 minutes. Once the supernatant was discarded, 70% ethanol was added to wash the DNA pellet. After the ethanol was discarded, the microcentrifuge tubes containing DNA samples were allowed to air-dry 20 minutes before being re-suspended in 50 µL of sterilized water for PCR. Concentrations of DNA samples were measured using a NanoDrop^TM^ 2000 Spectrophotometer (Thermo Scientific, USA).

### PCR analysis

Extracted genomic DNA from leaf tissues of putative transgenic lines and controls were used in PCR amplification for GUS gene. GUS gene (*gusA*)-specific primers **GUS-2**: 5’-CGTTTCGATGCGGTCACTCATTACG-3’ (forward primer) and **GUS-3**: 5’-TCTCCTGCCAGGCCAGAAGTTCTT-3’ (reverse primer) were designed and purchased from Invitrogen (USA). Promega Go*Taq*^®^ Flexi DNA polymerase kit was used for amplification. Each PCR reaction contained 3 µl (approximately 300 ng) of genomic DNA, 2 µl 2.5mM dNTPs, 2 µl 25mM MgCl_2_, 5 µl 5x PCR buffer, 1 µl of each primer (10 µM), 0.12 µl polymerase and water for a total volume of 25 µl. Thermocycle program used an initial denaturation at 94°C for 4 minutes, followed by 35 cycles of 94°C (30 seconds), 65°C (30 seconds) and 72°C (1 min 20 seconds), and a final extention step at 72°C for 2 minutes. All PCR were performed on an Eppendorf’s Mastercycler Gradient^®^ PCR machine (Eppendorf, USA). The PCR products were separated on a 1 % TAE agarose gel (Cat. No. 820723, MP Biomedicals, USA) stained with ethidium bromide (Cat. No. E3050, Technova, USA). The gel was photographed with a GelD°C-It™ Imaging System (Ultra-Violet Products LLC., USA).

### GUS histochemical assay

Putative transgenic lines and controls were tested for β-glucuronidase (GUS) expression according to Jefferson et. al. (1987). GUS was assayed by placing leaf tissues in the wells of a 96-well plate containing GUS-staining solution [1 mM 5-bromo-4-chloro-3-indoxyl-β-D-glucuronide (X-gluc)] (Gold BioTechnology, Inc., St. Louis, MO, USA), 100 mM sodium phosphate buffer pH7.0, 0.5 mM potassium ferricyanide, 0.5 mM potassium ferrocyanide, and 0.1% Triton X-100). After vacuum-filtration for 10 min, plate was incubated at 37°C overnight. To check GUS staining, chlorophyll of leaf tissue was removed by repeated washing in 70%. Chlorophyll interferes the observation of stained blue color. Stained leaf tissues were examined under a dissecting microscope and scored for blue coloration.

### Mendelian inheritance analysis of T_1_ seedlings

T_1_ seeds derived from kanamycin-resistant T_0_ putative transgenic lines were sterilized with ethanol and bleach, and then placed on the germination medium [MS mineral salts, 3% (w/v) sucrose, 1x Gamborg’s vitamin solution] supplemented with 100 µg/mL kanamycin and 200 µg/mL of mixture of cefotaxime and carbenicillin. Plates were placed in a growth chamber with 16-hr light/8-hr dark photoperiod. Antibiotic-resistant or susceptible plant seedlings were counted and documented for Mendelian inheritance analysis.

### Statistical analysis

The test of the “goodness of fit” of Mendelian ratio 3:1 (the ratio of resistant to susceptible seedlings) was carried out with the chi-square (χ^2^) test to estimate the number of single-locus transgenic lines at *p*=0.05 level.

## Results and Discussion

To check and produce material for transformation, seeds of four previously pN6.Bxb1-transformed lines were selected either on kanamycin- or hygromycin-containing medium. Transgenic ones should survive hygromycin selection, and die from kanamycin selection (Table 1). After 10-day selection, all seedlings died on <u>kanamycin</u>-containing plates (Fig. 3A), and resistant seedlings were observed on <u>hygromycin</u>-containing plates (Fig. 3B). The resistant plant seedlings grew normally with green cotyledons (and two tiny true leaves), having main roots growing and extending into the selection medium. The mature cotyledons are around 2 mm in diameter. In contrast, the growth of hygromycin-sensitive seedlings was stunted and have pale-green (or yellowish) cotyledons (Fig. 3B). The cotyledons were curvy and the roots could not grow and extend into the selection medium. Seedlings at the two-week-old stage were used for calculating the numbers and ratio of the resistant/susceptible seedlings to estimate transgene copy number (Table 1). Resistant seedlings were also used for secondary genetic transformation.

**Table 1.**
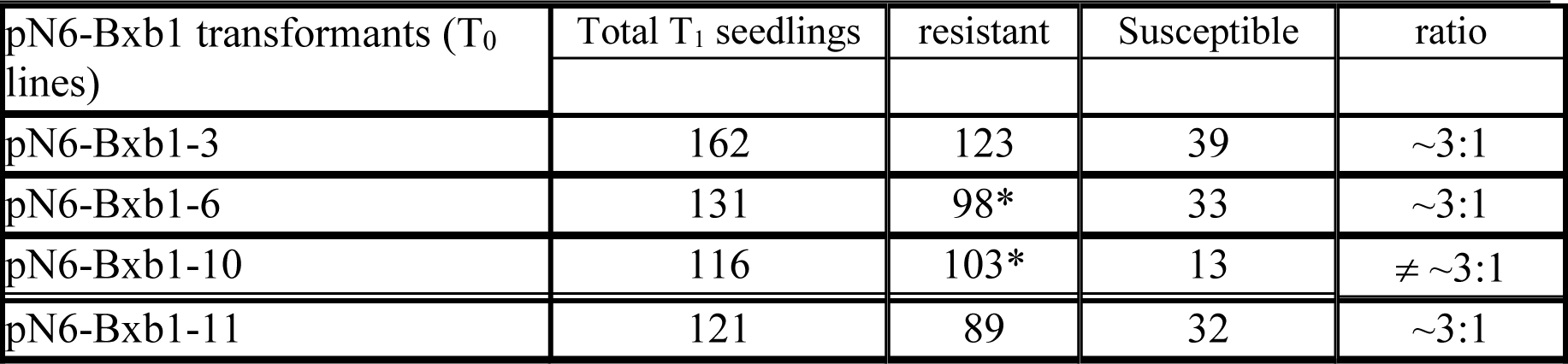
T_1_ seeds of pN6-Bxb1 transgenic lines were selected on MS medium containing antibiotic hygromycin. The survived plants were starting materials for secondary genetic transformation.

**Fig. 3.**
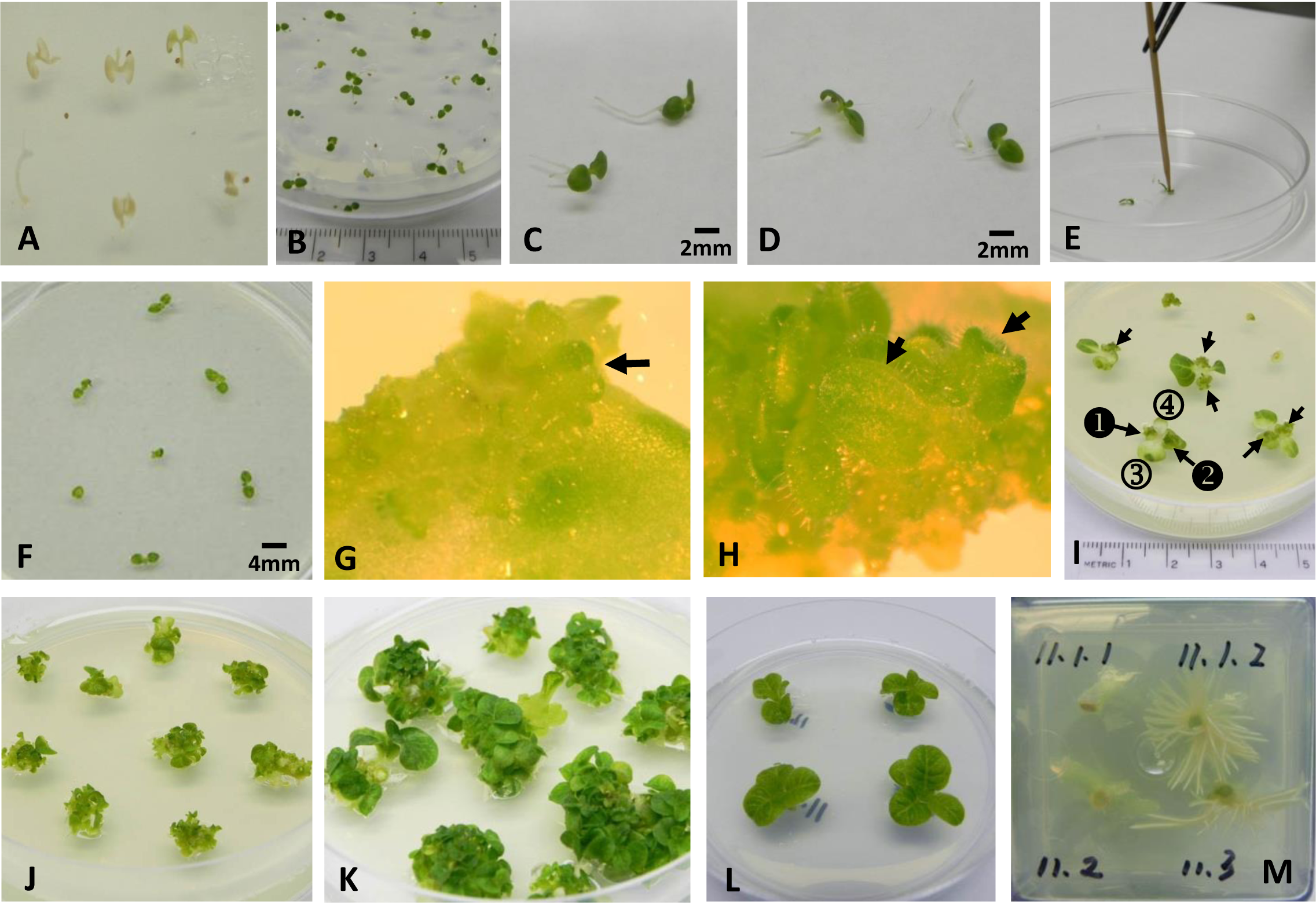
Toothpick inoculation of tobacco cotyledon. **A** Seedlings on kanamycin-containing MS plates. **B** Seedlings survived hygromycin selection. **C-D** Part of the main root was excised. **E** *Agrobacterium* inoculation with a sterile toothpick. **F** Two days of co-cultivation on MS medium, and then on selection medium. **G** Somatic embryoids observed 10 days after selection. **H** Germination of embryoids. **I-K** Putative transgenic shoots grew on the *Agrobacterium*-infected regions of explants (on selection medium). **L** Individual shoots were cut at the base and transferred to Rooting medium with selection agent. **M** Roots (plant on the right side) were observed in the selection medium.

Seedlings of 2-week-old surviving hygromycin selection were pulled out with a pair of forceps and placed in a sterile Petri-dish to excise the roots ~1.5 mm below the cotyledons (Fig. 2B and Fig. 3C-3D). Removal of most part of root eased transformation manipulation. The cotyledons were then used for *A. tumefaciens* strain LBA4404 (harboring pC35.BNK.2 or pRB140-Bxb1-op) transformation. Both pC35.BNK.2 and pRB140-Bxb1-op contain the *nptII* gene which confers resistance to antibiotic kanamycin for transgenic plants. Toothpicks were dipped in the *Agrobacterium* suspension and used to gently bruise the central area of the two cotyledons of the seedlings (Fig. 2C and Fig. 3E). After inoculation, the root-cut seedlings were placed on the selection medium abaxially, with the left root stuck into the medium (Fig. 3F). After 10 days of selection, embryoids were observed from the embryogenic calli around the wound area (Fig. 2D and Fig. 3G), and embryoid germination was seen at 2 weeks (Fig. 3H). A dissecting microscope is needed to observe the embryoids and their germination. The pair of true leaves (Fig. 3I ③ and ④) grew and expanded rapidly. The sizes became bigger than the cotyledons over days (Fig. 3I ❶ and ❷). However, under kanamycin selection, the expanded true leaves turned yellowish in color while the cotyledons with putative transgenic shoots remained green (Fig. 2E and Fig. 3I). The putative transgenic shoots may provide kanamycin-detoxified proteins to cross-protect the rest of the cotyledons and contribute to the green color of the cotyledons bearing these shoots. Cotyledons with putative transgenic shoots (Fig. 3I ❶ and ❷) were excised from the leftover-stem and placed in fresh selection medium to allow those putative transgenic shoots to grow further (Fig. 3J-3K). One to two putative transgenic shoots (several of them on each infected cotyledon) were then chosen and cut at the base after they reached 1 cm, and placed in the Rooting medium (with 100 µg/mL kanamycin) (Fig. 3L). To ensure the two putative transgenic shoots excised from each infected cotyledon were two independent transformation events, two shoots separated by a greater distance were carefully chosen. Roots could be seen within a week. Rooted plants around 5 cm in height were then transferred to soil for flowering and seed setting (Fig. 3M).

For pC35.BNK.2 transformation, a total of 56 putative transgenic shoots were excised from the infected cotyledons and allowed to grow in Rooting medium with selection. Among the 56 putative transgenic shoots, 38 of them (68%) grew roots in 100 µg/mL kanamycin-containing medium. 32 plants were randomly chosen and transplanted to soil. Among them, 28 of the plants survived transplanting and grew into adult plants. Except one plant presented with stunted growth, the other 27 plants showed wild-type architecture, were fertile and set T_1_ seeds. T_1_ seeds derived from 21 T_0_ transgenic plants were randomly chosen and plated out on kanamycin-containing MS medium. Among them, 17 lines showed resistance/susceptible segregation for kanamycin selection (Fig. 4B-4C), while wild-type seedlings were dead (Fig. 4A). Three lines [pN6.Bxb1 #3 (15.2), pN6.Bxb1 #11 (7.1) and pN6.Bxb1 #11(10)] had no surviving seedlings from kanamycin selection, indicating that their parental T_0_ lines were either selection escapees or gene-silenced transgenic plants (Table 2). T_1_ seeds of line pN6.Bxb1#11 (18) did not germinate. Using the data of kanamycin selection on the T_1_ seed from 21 putative transgenic plants, the chi-square (χ^2^) ‘goodness of fit test’ for 3:1 ratio (resistant: susceptible) were performed at *p*-value = 0.05 level. The results indicated that six individual lines should have single-locus transgene integration (indicated “3:1” in Table 2), while the remaining lines showed multiple-locus transgene integration (Table 2).

**Table 2.** Effects of kanamycin (100 µg/ml) on root growth of putative T_0_ transgenic lines (in Rooting medium) and T_1_ seeds (in Germination medium) from pC35.BNK.2 transformation. One or two T_0_ putative transgenic lines were randomly chosen from *Agrobacterium*-mediated cotyledon transformation for analysis. Segregation (resistant vs. susceptible to kanamycin) ratio of T_1_ generation on kanamycin-containing media was also analyzed. Goodness-of-fit for 3:1 ratio was determined using chi-square (χ^2^) test, at *p-value* = 0.05 level.

| Parental lines | T<sub>0</sub> putative transgenic lines | Rooting in kan medium (T<sub>0</sub>) | Kanamycin selection (T<sub>1</sub> seeding R:S ratio) |
| --- | --- | --- | --- |
| **pN6.Bxb1 #3** |  |  |  |
|  | 5 | +<sup>1</sup> (seeds)<sup>2</sup> | N/A<sup>3</sup> |
|  | 8.2 | + (seeds) | **<u>83:30 (3:1)</u>** |
|  | 9 | + (seeds) | 68:11 |
|  | 11 | + (seeds) | 130:9 |
|  | 12.1 | + [lost of plant] | N/A |
|  | 12.2 | + (seeds) | **<u>72:25 (3:1)</u>** |
|  | 15.2 | + (seeds) | 0:114 |
|  | 20 | + (seeds) | 12:0 |
|  | 21 | + (seeds) | 100:3 |
| **pN6.Bxb1 #11** |  |  |  |
|  | 1.2 | + (seeds) | 68:4 |
|  | 3 | + (seeds) | 118:14 |
|  | 5 | + (seeds) | **<u>35:13 (3:1)</u>** |
|  | 7.1 | + (seeds) | 0:66 |
|  | 7.2 | + (seeds) | N/A |
|  | 8.1 | + (seeds) | 103:21 |
|  | 8.2 | + (seeds) | **<u>91:30 (3:1)</u>** |
|  | 9 | + (seeds) | 107:18 |
|  | 10 | + (seeds) | 0:111 |
|  | 11 | + (seeds) | N/A |
|  | 13 | + (seeds) | 52:6 |
|  | 14 | + (seeds) | 36:7 |
|  | 15 | + (seeds) | N/A |
|  | 16 | + (seeds) | **<u>110:34 (3:1)</u>** |
|  | 17 | + (seeds) | **<u>99:29 (3:1)</u>** |
|  | 18 | + (seeds) | no germination |
|  | 19 | + (seeds) | 69:15 |
|  | 20 | + (seeds) | N/A |
|  | 21 | + (seeds) | N/A |

<sup>1</sup> ‘+’ indicates that roots grew into kanamycin medium
<sup>2</sup> Indicates that T<sub>0</sub> plants set T<sub>1</sub> seeds
<sup>3</sup> N/A: not applicable

**Fig. 4.**
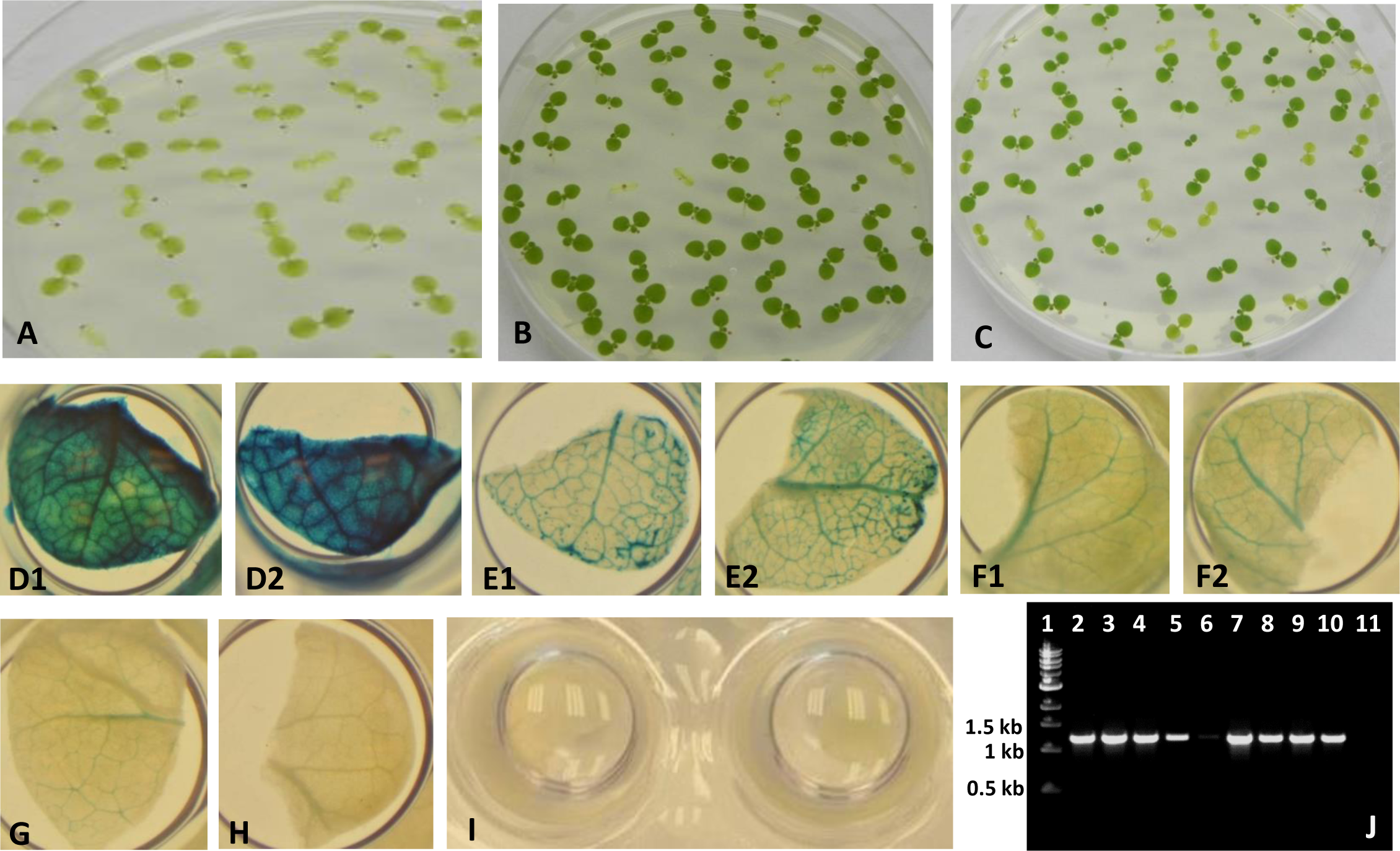
Mendelian inheritance analysis of T_1_ seeds from putative T_0_ transgenic lines (pC35.BNK.2 transformants) on kanamycin medium (A-C), and GUS-staining patterns of T_0_ transgenic lines from pRB140-Bxb1-op transformation (D-J). **A** Kanamycin selection: wild-type seedlings. **B-C** T_1_ seedlings of putatively pC35.BNK.2-transformed T_0_ lines. **D-H** GUS staining of leaf tissues from putatively pRB140-Bxb1-op-transformed transgenic lines. **I** GUS staining of wild-type plants. **J** PCR results of GUS gene from pRB140-Bxb1-op-transformed putative transgenic lines. Lane 1: DNA size markers, lanes 2-9: individual transgenic plants, lane 10: positive control, lane 11: water (negative control)

In summary, in this transformation study, stable transgenic lines can be obtained by using sterile toothpicks as a tool to deliver *Agrobacterium*. 81% (17 out of 21) T_0_ transgenic lines, which were randomly chose for analysis, showed stable insertion of *nptII* transgene and conferred kanamycin resistance in the pC35.Bxb1.2-transformation experiment. 35% (6 out of the 17) of the stable transformants demonstrated single-locus transgene integration deduced from chi-square (χ^2^) test. Transgenic plants with single-copy transgene insertions are preferred over those having multiple transgene copies, because the latter is prone to gene silencing (Tang et al. 2007). It was reported that the frequency of single-copy transgene insertion in *Arabidopsis* for *Agrobacterium*-mediated transformation was 15% (De Paepe et. al. 2009). Although demonstrated in a different species, this method has generated a higher percentage (35%) of single-locus transgene insertion transformants.

For pRB140-Bxb1-op transformation, using water-prepared *Agrobacterium* inoculum, we have also obtained putative transgenic lines (Table 3), at least 6 independent lines with dark-blue GUS-staining on their leaf tissues (representatives of 2 plants presented in Fig. 4 D1-D2). Other GUS-staining patterns were also observed (Fig. 4 E1-E2, F1-F2, G and H). Leaf tissue of wild-type did not stain blue (Fig. 4I). PCR of GUS gene amplification of those plant tissues showed expected size (Fig. 4J). The data indicated that transgenic plants can be produced using *Agrobacterium* suspension prepared with only water (*Agrobacterium* colonies re-suspend in water). We did not track these plants into adulthood. The purpose of this experiment was just to see whether transgene (*gus*) could be stably transformed and expressed using toothpick inoculation. Water-prepared *Agrobacterium* inoculum also successfully generated stable transgenic plants. This can save labor of medium preparation and funds for cost of medium. However, transformation efficiency between using medium-prepared or water-prepared *Agrobacterium* suspension was not compared in this study. This would require further experiments.

**Table 3.** Results from GUS-staining of leaf tissues from putatively pRB140-Bxb1-op-transformed individual transgenic lines. Water-prepared *Agrobacterium* culture used for transformation.

| Transgenic lines | GUS staining | Tissue with staining<sup>1</sup> |
| --- | --- | --- |
| 1 | **dark blue** | **all tissue [Fig.4D]<sup>2</sup>** |
| 2 | **strong blue** | **veins [Fig.4E]** |
| 3 | **strong blue** | **veins [Fig.4E]** |
| 4 | no blue staining appear | all tissue [Fig.4H] |
| 5 | very faint light blue | veins [Fig.4H] |
| 6 | light blue | veins [Fig.4F] |
| 7 | light blue | veins [Fig.4F] |
| 8 | light blue | veins [Fig.4F] |
| 9 | **dark blue** | **all tissue [Fig.4D]** |
| 10 | **dark blue** | **all tissue [Fig.4D]** |
| 11 | very faint light blue | veins [Fig.4H] |
| 12 | light blue | veins [Fig.4F] |
| 13 | light blue | veins [Fig.4F] |
| 14 | **strong blue** | **veins [Fig.4E]** |
| 15 | very light blue | veins [Fig.4G] |
| 16 | N/A<sup>3</sup> | N/A |
| 17 | very light blue | veins [Fig.4G] |
| 18 | N/A<sup>3</sup> | N/A |
| 19 | light blue | veins [Fig.4F] |
| Positive control<sup>4</sup> | Blue | transgenic seeds |

<sup>1</sup> Types of tissues stained blue
<sup>2</sup> Refer to photograph showed in Fig. 4
<sup>3</sup> No putative transgenic plants produced from cotyledons of these seedlings. Callus formed
<sup>4</sup> Previous GUS-gene transformed T<sub>1</sub> seeds (Yau et al. 2011) were used for positive control

No *Agrobacterium* overgrowth was observed in these experiments. Overgrowth is one of the major problems of plant genetic transformation, and *Agrobacterium* can be seen to grow out of control on explants and eventually destroy the explants (Liu et al. 2016). How to control *Agrobacterium* overgrowth is a frequently asked question in *ResearchGate*, the largest professional network for scientists (https://www.researchgate.net/). In most cases, once overgrowth occurs, it is impossible to reverse. The best known solution is to begin again with another transformation experiment. Instead of submerging the whole leaf-disk in *Agrobacterium* culture, this protocol uses *Agrobacterium* inoculation only on a small area of the cotyledons. This practice could potentially minimize the *Agrobacterium* overgrowth problem. There is no requirement for washing the infected explant (cotyledons) in antibiotic solution.

Submersion of leaf explants in liquids can also have negative impact on the fitness of tissues for later use in culture. Successful genetic transformation using explant ‘cut edge’ for *Agrobacterium* inoculation have also been reported in cotton (Sunikumar and Rathore 2001; Yau 2017). Although toothpick inoculation had been used for screening *Agrobacterium* clones on 4-8 month old tobacco true leaves for VIGS study (<u>https://www.plantsci.cam.ac.uk/research/davidbaulcombe/methods/vigs</u>), to our best knowledge, this is the first report of using toothpick inoculation for tobacco cotyledon transformation.

A species as well reported as tobacco, we did not perform Southern assay for these studies. Instead, stable gene expression (ex. from *nptII* and *gus*) results were the focus in this report.

## Conclusions

Toothpick inoculation method has demonstrated positive results with using both medium-based and water-based *Agrobacterium* suspension for the purpose of infecting of very young tobacco seedlings. Transgenic plants resistant to kanamycin were obtained by using sterile toothpicks to inoculate *Agrobacterium* on cotyledons. T_1_ generation of stable transgenic lines segregated for kanamycin-resistance and susceptibility were observed from pC35.BNK.2 transformation. GUS-positive transgenic lines obtained from pRB140-Bxb1-op transformation were also observed. Previous leaf-disk-transformation method involving the use of the true leaves of tobacco plants requires waiting on seedlings to develop leaves of adequate size for traditional leaf-disk transformation. This study demonstrates waiting for true leaves is not required for performing genetic-transformation of tobacco. This can help researchers save a minimum of several weeks wait time, by removing the need to wait for true leaves to be obtained from seeds. Also of note, when hygromycin-resistance is a characteristic of transformed seeds, researchers must wait an extended period of time for true leaves to appear, since seedlings positive for selection are weakened in selection media. Growth of seedlings positive for selection will be slow, increasing the time necessary for true leaves to appear, even after transfer to media free of antibiotics. This innovative method involving sterile toothpicks allows researchers to perform genetic transformation on the cotyledons of tobacco seedlings. A vital benefit to this technique is the reduction of wait time in the production of genetically-transformed tobacco plants. By avoiding the step of submerging the explants in *Agrobacterium* suspension can also minimize the occurrence of *Agrobacterium* overgrowth. The explants can also be better maintaining fitness for later tissue culturing.

## Acknowledgments

This work was partially supported by Northeastern State University Faculty Research Committee Grant P120000. The authors are grateful to Dr. Kevin Wang for assistance in this work. The authors also wish to thank Dr. C. Neal Stewart, Jr. for useful discussions and suggestions.

## Authors’ contributions

YYY designed the experiment, constructed the plasmids, collected data and interpreted the research results. YYY supervised ME and LB, prepared and submitted the manuscript. ME and LB provided technical assistance with plant tissue culture, medium preparation, sample collection, PCR and GUS analysis. ME also participated with manuscript preparation and editing.

